# Unaware processing of tools in the neural system for object-directed action representation

**DOI:** 10.1101/119081

**Authors:** Marco Tettamanti, Francesca Conca, Andrea Falini, Daniela Perani

## Abstract

The hypothesis that the brain constitutively encodes observed manipulable objects for the actions they afford is still debated. Yet, crucial supporting evidence demonstrating that such visuo-motor embodiment occurs even without awareness has hitherto not been provided. In this fMRI study, we reliably instantiated unaware visual perception conditions by means of Continuous Flash Suppression, and found consistent activation in the target visuo-motor, action representation system, specifically for manipulable versus non-manipulable objects

## Introduction

The neural network involved in the encoding, storage, and retrieval of manipulable object knowledge comprises left-lateralized premotor, parietal, and posterior temporal cortices^1-4^. Visual perception of manipulable objects activates this visuo-motor network, despite the absence of motor task requests^5-7^. However, it is debated whether visuo-motor coding is triggered by the mere object’s visual appearance, as a constitutive component of its embodied representation, or instead by ancillary action imagery and planning processes engendered by the visual awareness of an object^8-11^. An unexplored resolutive approach for this controversy is to test whether the visuo-motor, object-directed action representation system (OAS) is also activated by manipulable entities under unaware visual processing conditions. Available, indecisive evidence is limited to an involvement of posterior visual and visuospatial areas^12,13^. The current fMRI study was aimed at evaluating the recruitment of the OAS during the processing of subliminally^14^ presented pictures of manipulable objects, with specific neuroanatomical predictions based on a meta-analysis^2^ on tool-related cognitive representations (Online Methods, Supplementary Table 1).

We determined the subjective perceptual threshold (PT) in 24 healthy subjects, with a continuous flash suppression (CFS) paradigm^15^, requiring subjective rating along a perceptual awareness scale (PAS)^16^. During fMRI, we then used the same paradigm but with a new set of pictures reflecting Manipulability (manipulable objects: MO; non-manipulable objects: NO) and Contrast (5 incremental levels: 2 below PT, 1 at PT, 2 above PT). Importantly, we included a null-stimulus reference baseline as an objective control for the true absence of perception (Figure 1).

**Figure 1.**
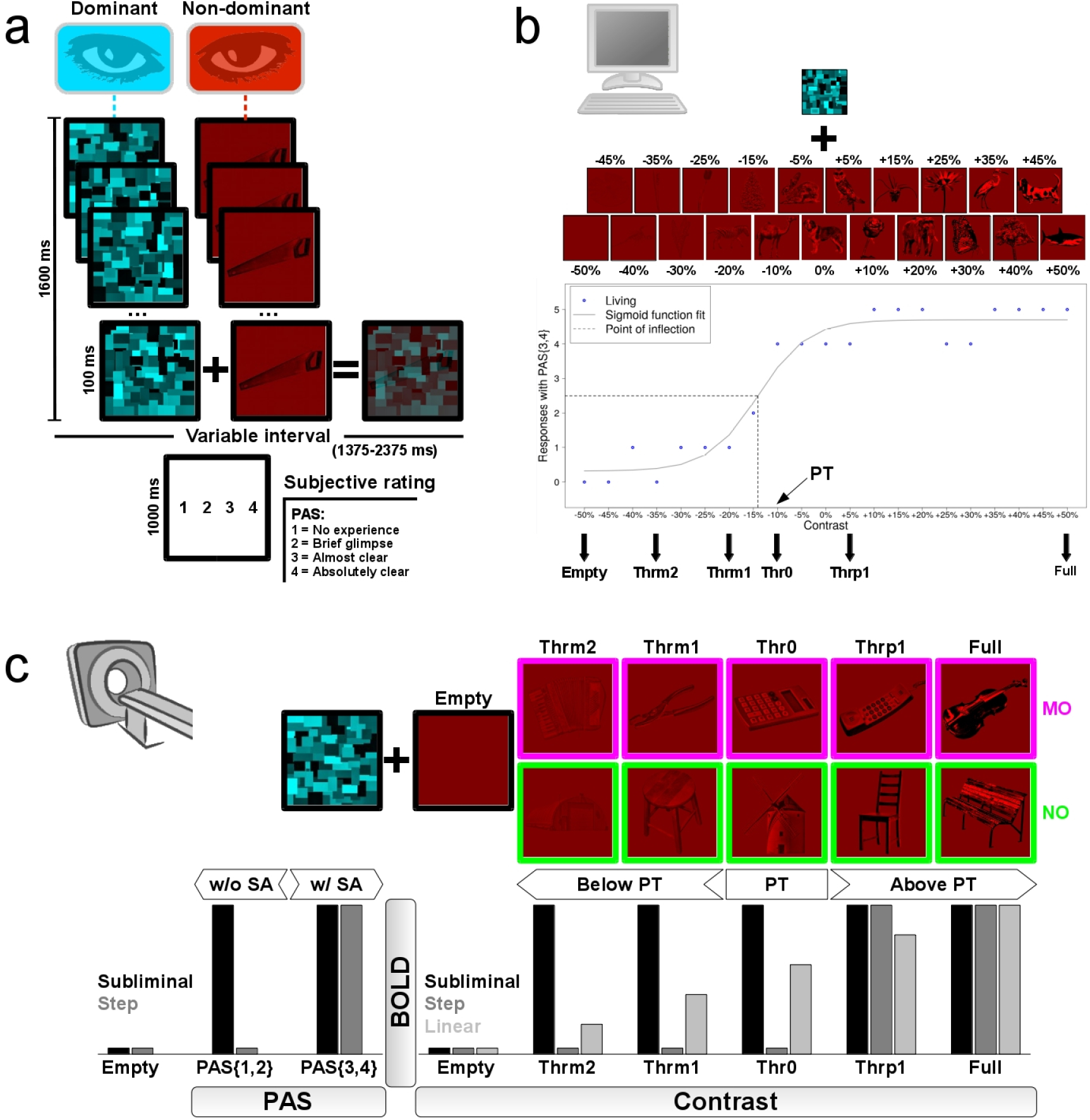
Experimental procedures. a) A single trial of the CFS task, with the presentation of a single picture at a given contrast level, overlaid by random texture masks flickering at 10 Hz. After a short interval, the participants reported a perceptual awareness scale (PAS) score, reflecting their stimulus’ subjective awareness (SA) level. The trial structure was identical for both the behavioral and fMRI sessions. b) In the pre-fMRI behavioral session, all masked stimuli belonged to the Living category. Each participant was presented with 5 trials for each of 21 image contrast levels, from full invisibility (-50%) to full visibility (+50%), with 5% increments. We plotted the number of responses (blue circles) with a PAS score of 3 or 4 (PAS{3,4}), indicating high to maximal SA. We then fitted a sigmoid function (Online Methods), and used its point of inflection to define the subjective perceptual threshold (PT). Six Contrast levels were then set to be used in the fMRI session: a void-of-object baseline (Empty), two contrast levels below PT (Thrm2, Thrm1), one at (Thr0), and two above (Thrp1, Full). c) In the fMRI session, the masked stimuli were either manipulable objects (MO) or non-manipulable objects (NO). We looked for MO-specific BOLD responses that fitted: bottom right) a “subliminal”, as opposed to a “step” or “linear”, profile of activation as a function of Contrast (i.e. activation both below and above PT); bottom left) a “subliminal” as opposed to a “step” profile as a function of PAS (i.e. both with and without SA).

## Results

We first analyzed the PAS behavioral responses during fMRI, to make sure that the image Contrast level significantly modulated the level of subjective awareness (SA). A generalized linear mixed model (GLMM) provided evidence that Contrast (Chi^2^(4) = 233.94, P <2.2 × 10^−16^), and not Manipulability (Chi^2^(1) = 0.38, P = 0.537), was a significant predictor of PAS ratings. The Contrast x Manipulability interaction was not significant (Chi^2^(4) = 0.98, P = 0.913): for both MO and NO, Contrast increase produced a nearly-linear increase of SA (Fig. 2a). Contrasts below the PT yielded low to minimal PAS scores, whereas those above the PT yielded high to maximal PAS scores.

**Figure 2.**
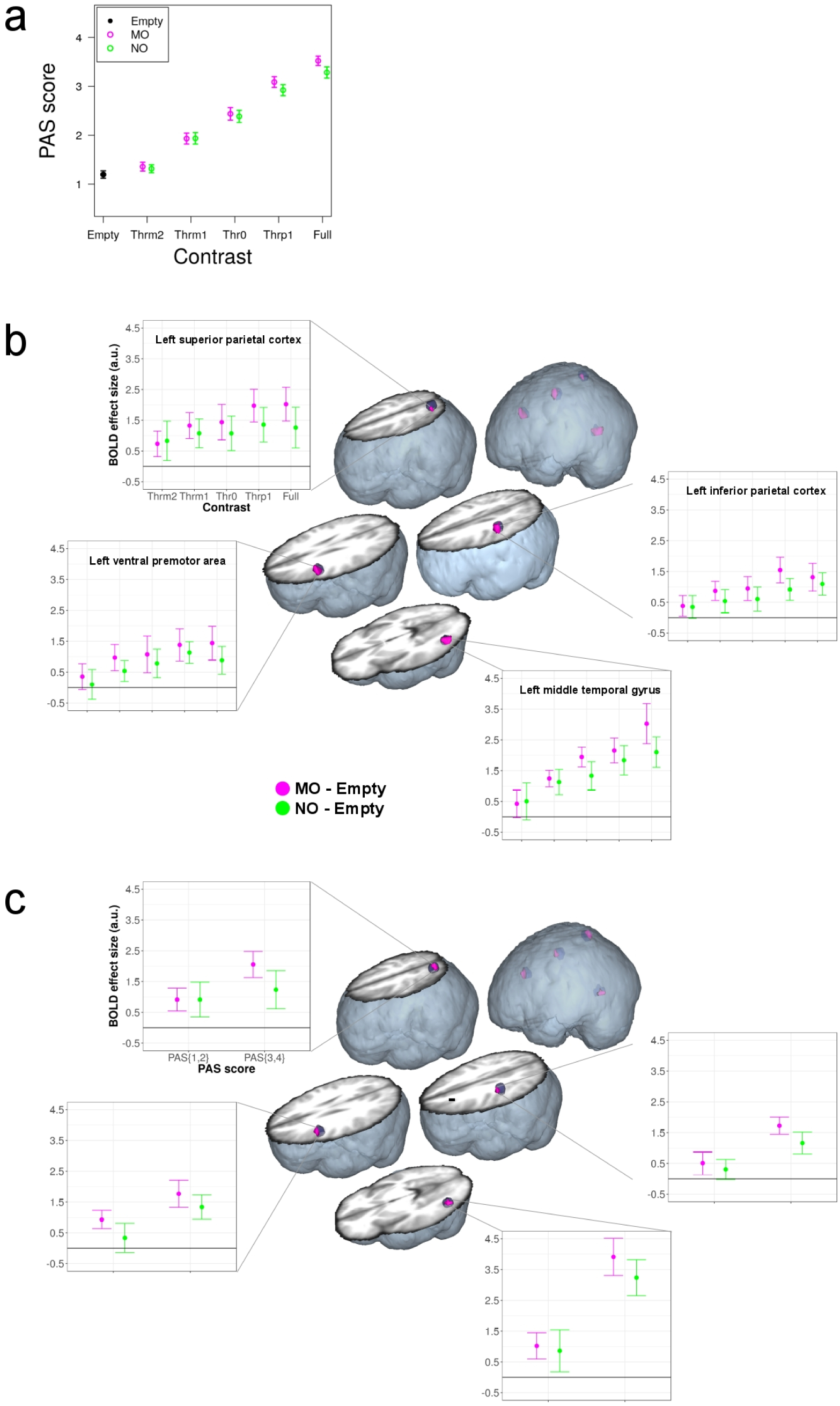
Behavioral and fMRI results. a) Average (n = 24) PAS ratings for MO and NO as a function of image Contrast level. Vertical bars indicate 95 % confidence intervals. b) Subliminal MO-specific brain activation in the OAS and in the left lateral middle temporal gyrus as a function of Contrast (n = 24). Brain activations (smFWE P < 0.05, purple color) are overlaid on the small volumes of interest (blue spheres) and displayed on a volumetric rendering of the average anatomical image of all participants. Dot-plots indicate average BOLD responses across all significant voxels in the activation cluster. Vertical bars indicate 95 *%* confidence intervals. c) Subliminal MO-specific brain activation in the OAS and in the left lateral middle temporal gyrus as a function of PAS ratings (n = 12, smFWE P < 0.05). All conventions are identical to panel b. PAS{1,2} = level of the PAS score factor corresponding to CFS trials in which the reported PAS score was either 1 or 2; PAS{3,4} = level for trials with reported PAS score 3 or 4.

As for neural activations induced by CFS, we first verified that, above PT, MO induced higher than NO response amplitudes in the OAS, as should be expected based on the visible visual features of each object category (Online methods, Supplementary Table 2a). We found stronger activation for MO versus NO in the ventral premotor, inferior parietal, and superior parietal cortices. Outside the OAS, in brain regions specifically involved in object identification (Supplementary Table 1), the lateral middle temporal gyrus also displayed stronger activation for MO versus NO, whereas the reverse effect (NO > MO) was found in the left and right fusiform giry (Supplementary Information, Supplementary Tables 3, 4).

Crucially, we sought for evidence that also below PT MO induced stronger than NO activation in the OAS. To this aim, we tested whether the levels of Contrast modulated OAS activation following either a “subliminal” (i.e. activation both above and below PT), a “step” (i.e. activation only above PT), or a “linear” (i.e. gradual activation increase with Contrast) BOLD amplitude model (Online Methods, Supplementary Table 2b). The strongest evidence indicated that MO-specific processing conformed to a subliminal model, in the ventral premotor area (small volume Family Wise Error (smFWE) type correction for multiple comparisons, P = 0.008, Z(1,23) = 3.36, 30 voxels, x = -50, y = 8, z = 28), inferior parietal cortex (smFWE P = 0.008, Z(1,23) = 3.35, 23 voxels, x = -48, y = -28, z = 44), and superior parietal cortex (smFWE P = 0.040, Z(1,23) = 2.73, 5 voxels, x = -28, y = -60, z = 60). MO-specific subliminal processing also extended into the lateral middle temporal gyrus (smFWE P = 0.004, Z(1,23) = 3.59, 37 voxels, x = -50, y = -68, z = -4) (Fig. 2b, Supplementary Fig. 1a, Supplementary Information, Supplementary Table 5).

To gain conclusive evidence that MO specifically activated the OAS, not only above but also under subliminal perceptual conditions, we further analyzed BOLD responses as a function of PAS ratings (Online Methods, Supplementary Fig. 2). This allowed us to test whether the presence of a MO stimulus elicited specific responses in the OAS also in the absence of SA, independently of image Contrast. The strongest evidence indicated that, as a function of PAS ratings, MO specifically activated the OAS conforming to a subliminal BOLD amplitude model (i.e. activation both with and without SA, Supplementary Table 6). Significant effects were located in the ventral premotor area (smFWE P = 0.013, pseudo t = 3.28, 13 voxels, x = -50, y = 12, z = 32), inferior parietal cortex (smFWE P = 0.032, pseudo t = 2.92, 8 voxels, x = -48, y = -28, z = 44), and in the superior parietal cortex (smFWE P = 0.017, pseudo t = 3.01, 10 voxels, x = -22, y = -66, z = 60). The lateral middle temporal gyrus (smFWE P = 0.022, pseudo t = 2.89, 12 voxels, x = -46, y = -66, z = -4) also presented a subliminal response (Fig. 2c, Supplementary Fig. 1b, Supplementary Information, Supplementary Table 7).

## Discussion

Resting upon unequivocal behavioral evidence of an absence of SA for object stimuli presented with image contrasts below PT, our fMRI results indicate that action-related properties of MO are capable of triggering a functional response in the left-lateralized premotor-parietal OAS and in the lateral posterior temporal cortex, even under unaware processing conditions. Among the brain regions displaying subliminal BOLD activation, the posterior temporal cortex showed the highest degree of linearity, with greater activation as Contrast and SA increased (Fig. 2b,c). Premotor-parietal cortices instead displayed a more stable activation level across image contrasts, below and above PT (Fig. 2b,c). Distinct response profiles in dorsal versus ventral visual areas as a function of awareness were also observed in previous studies^12,13^. Importantly, however, we show for the first time that both ventral and dorsal visual streams, extending into anterior parietal and premotor regions, support unaware processing of visual MO information. This is in good agreement with the observations in neurological conditions, where lesions selectively sparing either streams – the dorsal stream, e.g., in visual agnosia^17,18^; the ventral stream, e.g., in visuospatial neglect^19,20^ – can be associated to largely spared capacity to direct actions toward objects located in the surrounding space, despite lack of awareness.

Our results provide crucial evidence of the intimate neural coupling between visual perception and motor representation that underlies manipulable object processing: even when falling outside perceptual awareness, MO stimuli specifically engage the OAS. This coupling endows the brain with an efficient mechanism for monitoring and reacting to outside world’s stimuli escaping awareness.

## Acknowledgments

We thank Silvio Conte for help with MRI data acquisition, and Stefano F. Cappa and Marta Ghio for precious comment on the manuscript.

## Author contributions

M.T. and D.P. designed the experiment; M.T. and F.C. collected and analyzed the data; M.T., F.C., A.F., and D.P. organized and wrote the manuscript.

---

**Supplementary Figure 1.**
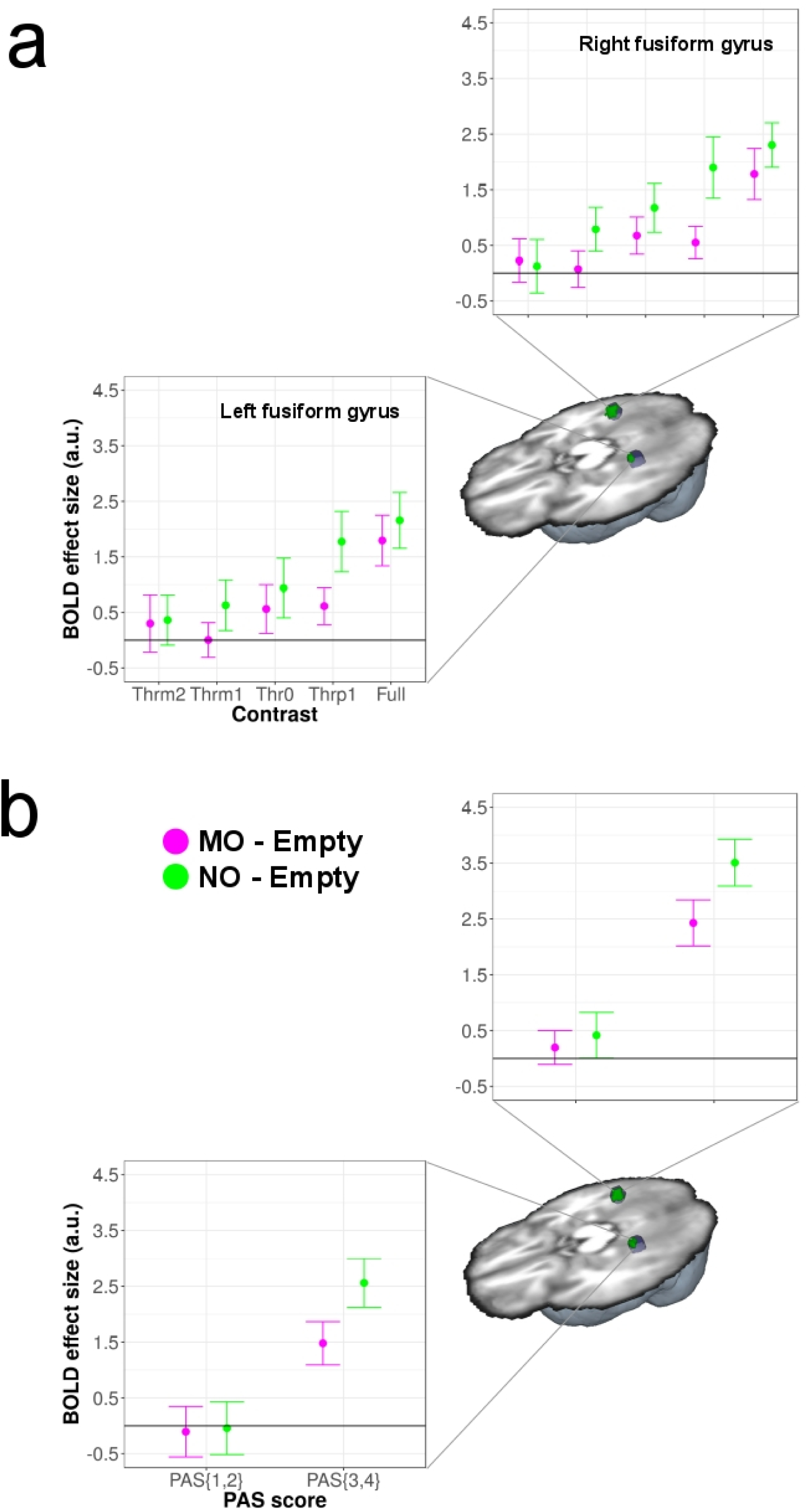
NO-specific brain activation in the fusiform gyri. a) Subliminal NO-specific brain activation as a function of Contrast (n = 24, smFWE P < 0.05), with significant effects in the left (smFWE P = 0.032, Z(1,23) = 2.82, 7 voxels, x = -26, y = -42, z = -12) and right (smFWE P = 0.001, Z(1,23) = 4.07, 18 voxels, x = 32, y = -46, z = -8) fusiform gyri. b) Subliminal NO-specific brain activation as a function of PAS ratings (n = 12, smFWE P < 0.05), with significant effects in the left (smFWE P = 0.016, pseudo t = 3.30, 10 voxels, x = -30, y = -42, z = -12) and right (smFWE P = 0.003, pseudo t = 5.01, 18 voxels, x = 30, y = -50, z = -8) fusiform gyri. All conventions are identical to Fig. 2 in the main text, except for the activations in green color to indicate NO-specific effects.

**Supplementary Figure 2.**
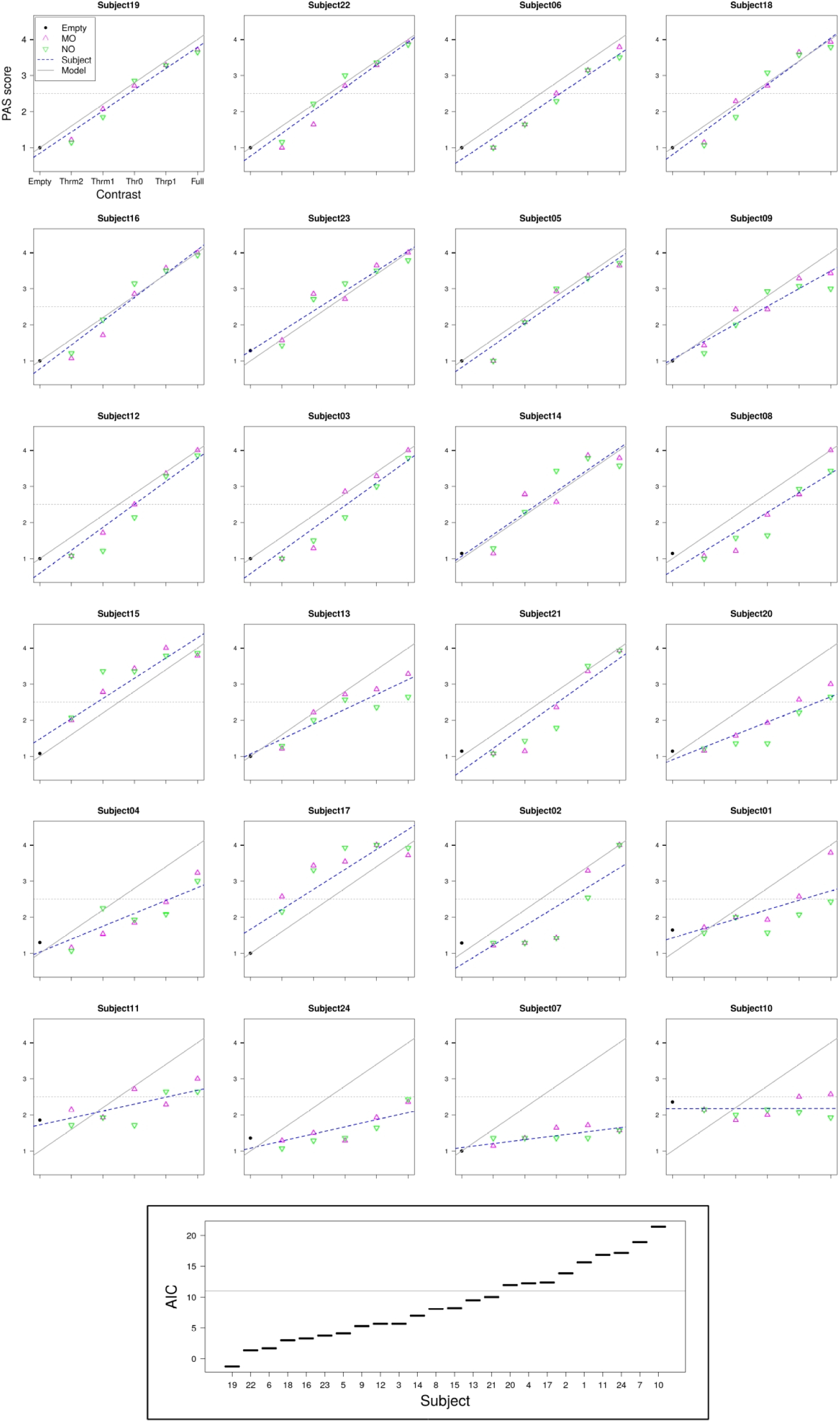
Selection of participants for the BOLD analysis as a function of PAS. This analysis required the inclusion of only those subjects that responded with comparable frequencies across all PAS levels (Online Methods). To this aim, the subjects were ranked according to the Akaike information criterion (AIC), obtained by Maximum Likelihood Estimation of the fit of the individual Contrast-to-PAS linear function (blue dashed line) to a Contrast-to-PAS linear model function (gray line). The sub-sample was defined by selecting the best-ranked subjects, until AIC discontinuity from the n-ranked to the n+1-ranked subject (horizontal line in bottom inset). Subjects 14, 15, and 23 were excluded from the sub-sample, since they presented higher than half-maximum (horizontal dashed line) PAS scores at contrast level Thrm1, indicating that a relatively high proportion of stimuli below PT were actually visible to these subjects. This left the sub-sample with 12 subjects. This sub-sample of subjects presented a comparable modulation of behavioral SA by image Contrast as the whole group (Supplementary Fig. 3a), as well as a comparable OAS activation as a function of Contrast both above and below PT (Supplementary Fig. 3b, Supplementary Table 8).

**Supplementary Figure 3.**
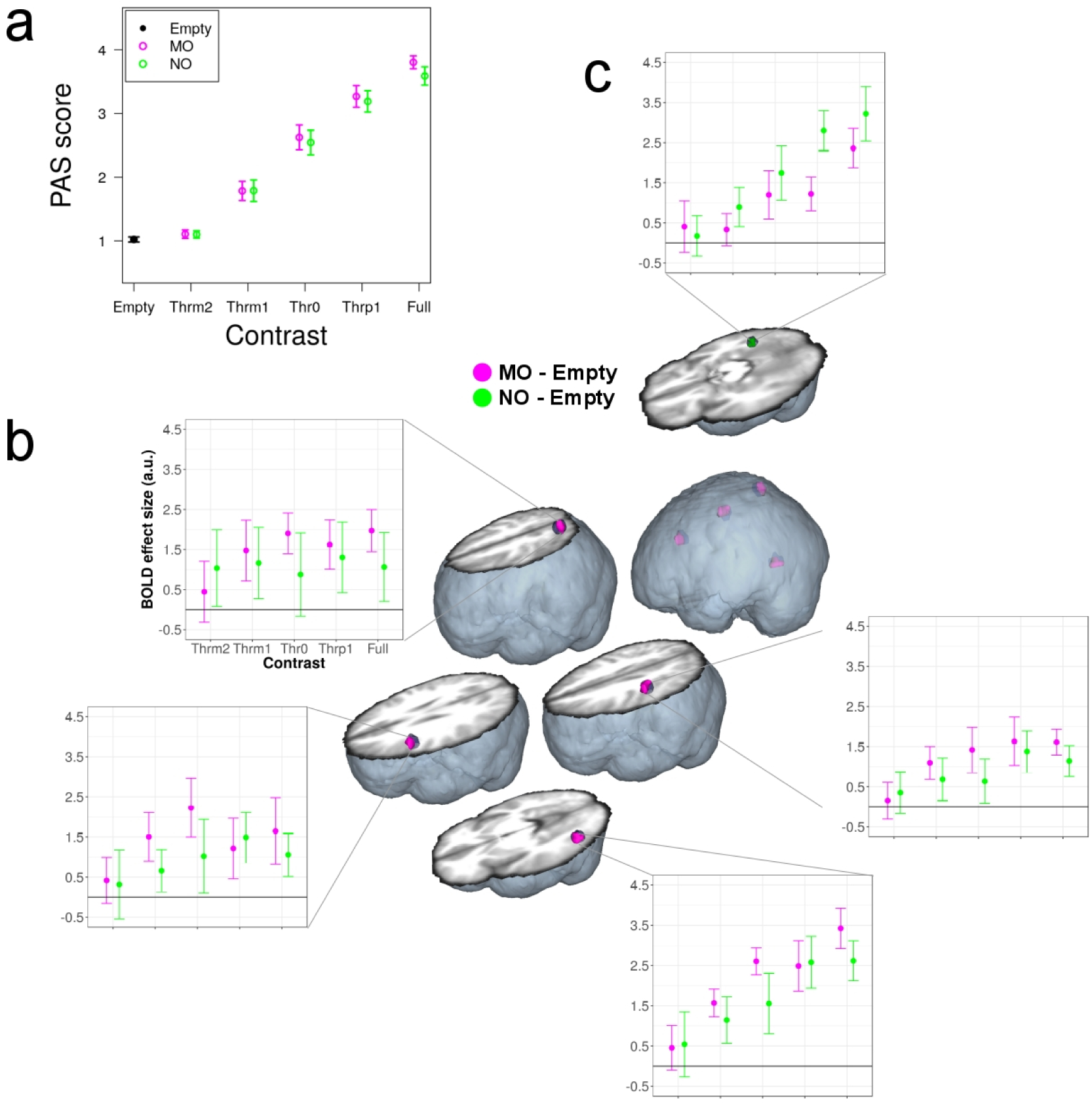
Behavioral and additional fMRI results in the sub-sample of 12 subjects included in the BOLD analysis as a function of PAS. a) Average (n = 12) PAS ratings for MO and NO as a function of image Contrast level. Vertical bars indicate 95 % confidence intervals. We found that Contrast (Chi^2^(4) = 248.51, P < 2.2 × 10^−16^), and not Manipulability (Chi^2^(1) = 0.15, P = 0.699), was a significant predictor of PAS ratings. The Contrast x Manipulability interaction was not significant (Chi^2^(4) = 0.46, P = 0.977). b) Subliminal MO-specific (purple color) brain activation as a function of Contrast (n = 12, smFWE P < 0.05). c) Subliminal NO-specific (green color) brain activation as a function of Contrast (n = 12, smFWE P < 0.05). All conventions are identical to Fig. 2 in the main text.

**Supplementary Figure 4.**
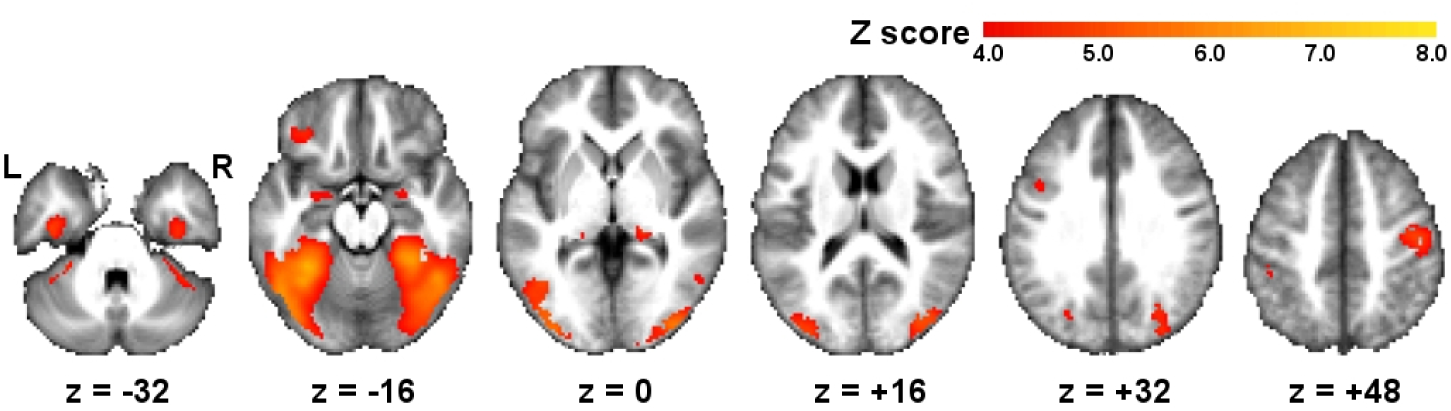
Main effect of Contrast (increases). Brain activations (n = 24, peak-level P < 0.05, whole brain FWE correction for multiple comparisons) are displayed in a red-yellow color scale reflecting the magnitude of Z scores, on axial sections (z coordinate level indicated in mm) of the average anatomical image of all participants. L = left; R = right.

## Online Methods

### Participants

Twenty-five Italian native speakers volunteered in the experiment. One participant did not comply with the task and was discarded from the analyses. All 24 included participants (13 females, mean age 22.09 years, SD = 2.19) were right-handed (mean score 0.95, SD = 0.07) according to the Edinburgh Inventory^21^. Eye dominance was evaluated for the CFS task (Fig. 1a) by means of the Miles test^22^: 4 participants were left, and 20 right eye dominant. All reported no history of neurological, psychiatric, or developmental diagnoses. They gave written consent to participate in the study after receiving a careful explanation of the procedures. The study was approved by the Ethics Committee of the San Raffaele Scientific Institute, Milano, Italy.

### Experimental stimuli

We used pictures of items belonging to three different semantic categories: manipulable objects (MO), non-manipulable objects (NO), and living entities. MO and NO pictures were employed for fMRI stimulation, living pictures for the pre-fMRI behavioral session.

All original pictures were high-resolution colored photographs, presenting a full shot of the depicted item in isolation, on a white background. The majority (85%) of pictures was drawn from the Bank of Standardized Stimuli (BOSS)^23,24^, whereas the remaining pictures (15%) were retrieved by internet search. MO (70 pictures) included objects whose specific function is carried out through manipulation with either one or both hands, such as utensils and musical instruments. NO (70 pictures) included objects whose specific function typically does not involve hand manipulation, such as buildings and seating furniture. Living entities (100 pictures) included animals and plants.

We ran a rating study on the set of pictures to collect rating norms for visual complexity^24^, and for two different manipulability measures^25^, that is, graspability and presence of functional motor associations. For each norming dimension, a distinct group of subjects rated all pictures on a 7-point Likert scale (1 = “low”, 7 = “high”). Visual complexity (5 rating subjects: 2 females, mean age 23.75 years, SD = 0.77) resulted balanced across the three semantic categories (MO: mean = 3.73, SD = 1.06; NO: mean = 3.84, SD = 1.10; Living: mean = 3.73, SD = 0.73; Kruskal Wallis Chi^2^(2) = 1.103, P = 0.576). In turn, the two manipulability measures resulted unbalanced, reflecting the specificity of MO, as opposed to NO and Living items: graspability (5 rating subjects: 3 females, mean age 23.52 years, SD = 0.27; MO: mean = 6.58, SD = 0.49; NO: mean = 2.80, SD = 1.38; Living: mean = 3.13, SD = 1.10; Kruskal Wallis Chi^2^(2) = 148.948, P < 0.001); functional motor associations (5 rating subjects: 4 females, mean age 24.30 years, SD = 0.78; MO: mean = 6.70, SD = 0.30; NO: mean = 2.39, SD = 1.07; Living: mean = 1.80, SD = 0.54; Kruskal Wallis Chi^2^(2) = 153.882, P < 0.001).

To make the stimuli suitable for the CFS task, we submitted all pictures to a customized image processing pipeline. First, all pictures were converted to black and white images, using ImageMagick 6.9 (www.imagemagick.org). Second, the mean brightness value of each picture was adjusted to the overall mean brightness value calculated over the entire picture set, using Matlab R2011a (MathWorks). Third, we increased and equalized the contrast of all pictures by means of Contrast Limited Adaptive Histogram Equalization (CLAHE)^26^ using Python 4.2 (www.python.org). Fourth, the image contrast was normalized across the entire picture set, again using Matlab R2011a.

With this normalized picture set, we then proceeded to generate the full gradation of image contrast levels required for our implementation of the CFS task (Fig. 1), which is based on the evidence – exploited for instance in the breaking-CFS task variant – that stimuli with higher contrast emerge more promptly from suppression^27^. For this purpose, we used GIMP 2.8 (www.gimp.org) to, firstly, replace the white image background with gray color (RGB values: 128, 128, 128), and, secondly, to progressively modify the object contrast by 5% incremental/decremental steps. This yielded, for each picture, a gradation of 21 different image contrast levels (Fig. 1b), ranging from -50% (i.e. full absence of object) to +50% (i.e. full visibility).

Finally, again using ImageMagick 6.9, we applied a red-channel hue to the full set of black and white images of 21 contrast levels, thus generating the final pictures that were presented to the non-dominant eye during CFS (Fig. 1a).

As for the masks presented to the dominant eye, we created a texture of small rectangles of different sizes and different gray color shades, chosen to roughly match the visual characteristics of the to-be-suppressed object pictures^27^. We randomized the position of the rectangles to generate 20 different mask exemplars. These were then colored with ImageMagick 6.9, by applying a cyan-channel hue (Fig. 1a).

### CFS task

The CFS task was organized as a series of consecutive trials, with trial structure identical for both the pre-fMRI behavioral and fMRI sessions (Fig. 1a). In both sessions, we used Presentation 18.3 (Neurobehavioral Systems) for stimulus delivery and behavioral response collection.

In each trial, a single object picture was presented for 1600 ms. During the same time interval, overlaid on this picture with 50% transparency (alpha blending), we presented a series of flickering masks at a frequency of 10 Hz (Fig. 1a). For each trial, 16 masks were randomly drawn, without replacement, from the pool of 20 available exemplars.

After a variable within-trial interval (randomized duration of 1375 ms, 1875 ms, or 2375 ms, in 4:2:1 proportion), the participants were presented for 1000 ms with the display “1 2 3 4”, and had to press one among four available buttons, according to their subjective visual perception of the stimulus. Before task administration, the participants were instructed and trained (by means of a few trials with explicit feedback from the experimenters) to associate each of the four buttons with a corresponding PAS^16^ rating level. The four rating levels indicated increasing levels of subjective stimulus perception, from lowest to highest: 1: no experience; 2: brief glimpse; 3: almost clear image; 4: absolutely clear image. PAS scores have been found to be highly predictive of the level of performance in visual identification of subliminal stimuli^28^, and to be concordant with objective measures of awareness^29^.

We intentionally avoided questions assessing objective stimulus perception, since these could bias visual object processing, by making the focus on the object’s semantic category or features explicit. Importantly, we nevertheless included a null contrast condition (void-of-object Empty baseline, Fig. 1) that constituted an objective control of absence of perception. The null contrast condition was used as a reference both in the behavioral and fMRI data analyses.

The between-trial intervals had a randomized duration of 2875 ms, 4125 ms, or 5125 ms (in 4:2:1 proportion).

Dichoptic stimulus view was instantiated by equipping the participants with red-cyan anaglyph plastic goggles^30^. The participants wore the cyan lens on the dominant, and the red lens on the non-dominant eye, thus establishing effective suppression of the red-hue object picture (Fig. 1a).

### Psychophysical assessment of PT

In a pre-fMRI behavioral session, performed on a laptop computer (Fig. 1b), every participant was presented with 105 consecutive CFS trials. Only pictures of Living entities were included in this session, in order to avoid pre-exposure with MO and NO pictures underlying the fundamental neuroimaging research questions.

Presentation 18.3 was coded such that, for each participant, the 100 Living available entities were randomly ordered in a list: the entities were then sequentially taken from the list, and assigned in batches of 5 to one of 20 image contrast levels (from -45% to +50%, Fig. 1b). This effectively instantiated a draw without replacement of each item in the available pool of 100 Living pictures, with an equal number of items for each contrast level. Additionally, 5 stimuli with -50% contrast level (void-of-object baseline) were also taken. The total number of 105 picture stimuli were then presented in semi-randomized order, each stimulus in a separate CFS trial.

For every participant, we plotted for each of the 21 image contrast levels the number of responses (possible range: 0-5, given 5 trials x contrast) with a PAS score equal or higher than 3 (PAS{3,4} score), indicating high to maximal SA of visual stimulus perception (Fig. 1b, blue circles). Using nonlinear least-squares estimation in Matlab R2011a, we then fitted this PAS{3,4} score and calculated the parameters (L, R, i, W) of a psychometric sigmoid function *f:*

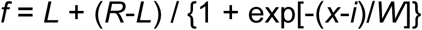
where: x = Contrast level; *L* = left horizontal asymptote (initial value == 0); *R* = right horizontal asymptote (initial value == 5); *i* = point of inflection (initial value == 2.5); *W* = width of the rising interval (initial value == 1). The estimated point of inflection *i* was taken as a reference for the subjective PT, defined as the first Contrast level above the point of inflection (Fig. 1b).

### fMRI Experimental design

In the fMRI CFS task (Fig. 1c), only MO and NO pictures were used. We factorially manipulated the two factors Manipulability (2 levels: MO, NO) and Contrast (5 levels: Thrm2, Thrm1, Thr0, Thrp1, Full). Four levels of the Contrast factor were individually tailored for each participant, based on the estimated subjective PT. Two of the tailored contrast levels were below threshold (Thrm2 = PT - 25%; Thrm1 = PT - 10%), providing subliminal stimulus presentation. One contrast level was at threshold (Thr0 = PT), and one was above threshold (Thrp1 = PT + 10%). The fifth contrast level was fixed for all participants as the maximum available contrast (Full). A null contrast control condition, namely a red square devoid of any objects, was also included for all participants (Empty, Fig. 1c). The Thrm2, Thrm1, Thr0, Thrp1, and Full contrast increments/decrements were chosen to roughly correspond to a logarithmic scale, on the basis of which we expected a roughly linear increase of PAS score responses (Fig. 1c)^13^. However, one participant had a PT corresponding to the -35% contrast level, and the contrast levels below threshold had thus to be tailored to a narrower range (Thrm2 = PT - 45%; Thrm1 = PT - 40%). Another participant had a PT corresponding to the +40% contrast level, and we therefore set the contrast level above threshold to a narrower range (Thrp1 = PT + 45%).

In the MRI scanner, the CFS task was carried out in two separate fMRI acquisition runs. For this reason, the available pool of 70 MO and 70 NO pictures was equally divided in two lists, each one containing 35 MO and 35 NO pictures. Presentation 18.3 was coded in such a way that, for each fMRI run, based on a semi-randomization of the list of 35 MO and 35 NO pictures that was invariant across participants, batches of 7 MO plus 7 NO pictures were drawn without replacement in sequential order and assigned to each Contrast level (Thrm2, Thrm1, Thr0, Thrp1, Full, in this exact order). Additionally, 7 stimuli with -50% contrast level (Empty baseline) were also taken. In this manner, all participants were presented with exactly the same depicted objects for every Contrast level, although the specific image contrast applied varied according to the individual PT (e.g., for level Thrm1, image contrast -20% for Subject03, and image contrast -5% for Subject04). For each of the two fMRI runs, the selected 77 picture stimuli were then presented in semi-randomized order, each one in a separate CFS trial.

Given this semi-randomization procedure, and the CFS trial structure described above (‘CFS task’ section), the stimulus presentation resulted in a fMRI event-related design, with semi-randomized stimulus presentation order (same randomization for all participants), and variable within-trial and between-trial intervals.

The order of the within-trial and between-trial intervals was determined by OPTseq2 (surfer.nmr.mgh.harvard.edu/optseq), in order to maximize the hemodynamic signal sensitivity of the event-related design.

Before the experimental fMRI runs, a brief fMRI training session was administered to each participant, in order to verify that the participant complied to the task instructions and requests.

### MRI data acquisition

Neuroimaging data were acquired on the same day of the psychophysical PT assessment. MRI scans were acquired with a 3 Tesla Philips Achieva whole body MR scanner (Philips Medical Systems, Best, The Netherlands) using an eight-channel Sense head coil (Sense reduction factor = 2). Whole-brain functional images were obtained with a T2*-weighted gradient-echo, EPI pulse sequence, using BOLD contrast (TR = 2000 ms, TE = 30 ms, flip angle 85°). Each functional image comprised 34 contiguous axial slices (4 mm thick), acquired in interleaved mode (field of view = 240 × 240 mm, matrix size = 128 × 128). Each participant underwent two consecutive functional scanning sessions, each comprising 313 scans, preceded by 5 dummy scans that were discarded prior to data analysis, and lasted 10 min and 36 s. A high-resolution T1-weighted anatomical scan (three-dimensional spoiled-gradient-recalled sequence, 1500 slices, TR = 7.2 ms, TE = 3.5 ms, slice thickness = 1 mm, in-plane resolution 1 × 1 mm) was acquired for each participant.

The stimuli were back-projected on a screen located in front of the scanner visible to the participants through a mirror placed on the head coil above their eyes. The participants gave PAS score responses through an MRI-compatible fiber-optic response box with 4 buttons, using their left hand. The left hand was chosen to reduce contamination of the BOLD signal measured in the target left-hemispheric OAS. As a further mean to make this contamination negligible, the variable within-trial interval introduced a temporal lag between the visual stimulus and the button press motor response, while also reducing the temporal correlation between the two events.

### GLMM analysis of behavioral data collected during fMRI

To evaluate the effects of Manipulability and Contrast on the PAS scores measured during fMRI, we run a nested series of generalized linear mixed models (GLMMs) using R 3.3.2^31^. We started with the simplest model in a nested hierarchy of increasingly complex GLMMs, with PAS scores as a categorical (4-point PAS scale) dependent variable, a fixed-effect modeling the 2 fMRI runs, and, as random intercept effects, the participants and the picture stimuli, to account for between-subjects and between-stimuli variability. The fixed-effects predictors of PAS scores were then added step-wise to increasingly complex GLMMs. The fixed-effects predictors were Manipulability (MO, NO), Contrast (Thrm2, Thrm1, Thr0, Thrp1, Full), and their 2-way interaction. All GLMMs were fit to a Poisson distribution and with a log-link function. Each hierarchically more complex model was tested against the hierarchically simpler model by means of a Chi^2^ log-likelihood ratio test (declared significance alpha-level: 0.05) to evaluate whether there was a significant increase in model fit, and thus a significant effect of the added predictor.

### fMRI data analysis

We used SPM12 (version 6685, www.fil.ion.ud.ac.uk/spm) for MRI data preprocessing and statistical analysis. The Segment procedure was applied to the structural MRI images of each participant, with registration to the Montreal Neurological Institute (MNI) standard space. Functional images were corrected for slice timing, and spatially realigned. The images were normalized to the MNI space, using the Segment procedure with the subject-specific segmented structural images as customized segmentation priors. Finally, the images were spatially smoothed with an 8-mm FWHM Gaussian kernel.

We adopted a two-stage random-effects statistical approach. The statistical analysis was restricted to an explicit mask including only the voxels with gray matter tissue probability > 0.1, based on the segmented structural images of each participant.

### Statistical analysis of fMRI activation as a function of Contrast (n = 24)

At the first stage, we specified a General Linear Model for each participant, with the time series high-pass filtered at 128 s and pre-whitened by means of an autoregressive model AR(1). No global normalization was performed. Hemodynamic evoked responses for all experimental conditions were modeled as canonical hemodynamic response functions. We modeled two separate sessions, each including 11 regressors of interest (Empty, MOThrm2, MOThrm1, MOThr0, MOThrp1, MOFull, NOThrm2, NOThrm1, NOThr0, NOThrp1, NOFull), with evoked responses aligned to the onset of each trial, and duration equal to the presentation of the masked stimuli (1600 ms). Separate regressors modeled experimental confounds, including PAS score responses, aligned to the onset of the “1 2 3 4” PAS display, task instructions, and head movement realignment parameters. If present, confound regressors were also specified for miss trials, and for trials with responses given before the appearance of the “1 2 3 4” PAS display, thus eliminating trials in which an anticipated motor response may contaminate the BOLD signal evoked by masked objects. Within the estimated first-level General Linear Model, we defined: i) a set of 10 condition-specific contrasts, each with a weight of +1 for the regressor of interest (e.g., MOThrm2) and a weight of -1 for the Empty baseline. ii) a set of 6 contrasts, modeling the variation of BOLD signal with the increase of image Contrast for MO and NO, according to either a subliminal, a step, or a linear BOLD amplitude model (Supplementary Table 2b).

Using the set of 10 condition-specific contrasts of each participant, we specified a second-level, random-effects, full factorial design. The model included the within-subjects factors Manipulability and Contrast (with the Empty baseline subtracted), with dependence and equal variance assumed between the levels of both factors. In this full factorial design, we investigated (Supplementary Table 2a) the simple effect of Manipulability for Contrast levels above PT, the main effect of Manipulability, and the main effect of Contrast (Supplementary Information).

Using the 6 Contrast-to-BOLD amplitude model contrasts of each participant, we specified three second-level, random-effects, paired t-test designs, investigating whether the activation differed between MO and NO as a function of image Contrast level following, respectively, a subliminal, a step, or a linear distribution profile. In each paired t-test design, the pairs included the two respective MO and NO contrasts of each participant (e.g., “MO subliminal model” and “NO subliminal model”, see Supplementary Table 2b). We assessed both MO-specific (MO > NO) and NO-specific (NO > MO) effects. For an unambiguous, one-tailed interpretation of these effects, which are otherwise confounded by the two-tailed nature of the first-level contrast images, we masked each second level effect with an inclusive binary mask given by the one-sample t-test statistics of the effect’s first term (e.g. “MO subliminal model > NO subliminal model”, inclusively masked by “MO subliminal model”).

### Statistical analysis of fMRI activation as a function of PAS ratings (n = 12)

Contrary to the analysis of BOLD responses as a function of Contrast, which rested on an equal number of CFS trials for each Contrast level, the analysis as a function of PAS ratings was complicated by unpredictable individual behavioral variability, with some participants markedly departing from a linear increase of PAS score with increasing Contrast level (Supplementary Fig. 2), thus producing an unbalanced number of trials across the 4 PAS scale levels. In order to minimize the problem of unbalanced statistical comparisons, the analysis was restricted to only those subjects that responded with comparable frequencies across all PAS levels. This amounted to reducing the sample of participants to a sub-sample (n = 12), by estimating with R 3.3.2 the maximum likelihood of the individual Contrast-to-PAS linear function to a Contrast-to-PAS linear model function, and then by ranking the participants based on the Akaike information criterion (Supplementary Fig. 2).

For this sub-sample of 12 participants, we specified at the first stage a General Linear Model, with all parameters identical to the one modeling BOLD responses as a function of Contrast, but this time reassigning the CFS trial events based on the PAS scores. This resulted in modeling 5 regressors of interest: the Empty baseline (unmodified); MOPAS{1,2}: all trials with MO pictures in which the reported PAS score was either 1 or 2; MOPAS{3,4}: all MO trials with PAS score 3 or 4; NOPAS{1,2}: all NO trials with PAS score 1 or 2; NOPAS{3,4}: all NO trials with PAS score 3 or 4. Within the estimated first-level General Linear Model, we defined a set of 4 contrasts, modeling the variation of BOLD signal with the increase of PAS score for MO and NO, according to either a subliminal or a step BOLD amplitude model (Supplementary Table 6).

Given the small sample size (n = 12), all second-level, random-effects analyses of BOLD activation as a function of PAS ratings were carried out using non-parametric statistics, by mean of the SnPM13 toolbox^32^. For each small volume of interest (6-mm radius spheres centered on the coordinates indicated in Supplementary Table 1), we specified a set of 2 paired t-test models, investigating whether the activation differed between MO and NO as a function of PAS score following, respectively, a subliminal or a step distribution profile. In each paired t-test design, the pairs included the two respective MO and NO amplitude model contrasts of each participant (e.g., “MO subliminal model” and “NO subliminal model”, see Supplementary Table 6). We assessed both MO-specific (MO > NO) and NO-specific (NO > MO) effects. Also for the non-parametric t-test analyses, we disambiguated the directionality of the effects by means of an inclusive binary mask given by the non-parametric one-sample t-test statistics of the effect’s first term (e.g. “MO subliminal model > NO subliminal model”, inclusively masked by “MO subliminal model”).

### fMRI analyses: declared significance threshold

For all the reported fMRI analyses, the significance threshold was declared at peak-level P < 0.05, using a small volume Family Wise Error (smFWE) type correction for multiple comparisons, with spherical small volumes of 6-mm radius centered on the coordinates of interest (Supplementary Table 1). The only exception was the main effect of Contrast (Supplementary Information), which was orthogonal to the Manipulability factor and related hypotheses, and for which we declared a peak-level P < 0.05 threshold, FWE-corrected at the whole brain level.

### Neuroanatomical predictions for tool-related cognitive representations

In the meta-analysis by Ishibashi et al.^2^, a cognitive fractionation is proposed, distinguishing between those functional processes related to planning and executing actions directed towards tools, and those related to tool identification. The latter, implicated in tasks calling for recognition and naming of objects, were shown by the meta-analysis to specifically activate the fusiform gyrus, bilaterally, and the left lateral occipito-temporal cortex. The former, implicated in planning, imagination, and execution of object manipulation, were instead specifically associated to activations in the left dorsal premotor and superior parietal cortices. Both identification and action processes were shown to conjointly activate the left ventral premotor and inferior parietal cortices.

In our study, we relied on the full set of these 7 either process-specific or shared brain regions for the definition of neuroanatomically constrained small volumes of interest (Supplementary Table 1). We differentiated the brain regions that are involved – either exclusively or in conjunction with identification – in object-directed action representation (i.e. left ventral and dorsal premotor, and inferior and superior parietal cortices), from the brain regions involved in object identification alone (i.e. bilateral fusiform gyrus, and left occipito-temporal cortex). The object-directed action representation system (OAS) constituted the core of our testing hypotheses.

The predictions on the brain regions specifically involved in object identification were instead more nuanced, in relation to the fundamental comparison carried out in our study between manipulable (MO) and non-manipulable (NO) objects. A close inspection of the control conditions employed in the studies entering the meta-analysis^2^ showed that, while there is indeed strong evidence of an activation of the inferior occipito-temporal cortex for tool identification, this activation has been typically evidenced by subtraction with low-level baseline conditions, such as rest, fixation, or scrambled object pictures. Only a very limited number of studies used subtraction baseline conditions that can be assimilated to NO, such as vehicles or seating furniture. Therefore, there is no strong principled evidence that allows to confidently predict stronger activation in the inferior occipito-temporal cortex for MO versus NO identification. On the contrary, a number of other fMRI studies has shown specific activation, particularly in the bilateral fusiform gyrus, for objects that are not associated with manipulation, such as vehicles and buildings^33,34^. Our predictions on the object identification system, and particularly on the fusiform gyri, were therefore of either no activation differences for MO and NO, or stronger activation for NO versus MO.

### Data availability

The data that support the findings of this study are available from the corresponding author upon request.

## Supplementary Information

### fMRI activation as a function of Contrast (n = 24): simple effect of Manipulability for image Contrast levels above PT (Thrp1, Full), and main effect of Manipulability

The NO-specific BOLD responses that we found in the fusiform gyri (Supplementary Tables 3, 4) are consistent with the latter’s involvement in the representation of non-manipulable inanimate objects^33,34^.

### fMRI activation as a function of Contrast (n = 24), conforming to step and linear BOLD amplitude models

In the OAS, there was no evidence of MO-specific step BOLD modulation. NO-specific step BOLD modulation was found in the bilateral fusiform gyri, with a concomitant linear activation profile in the right fusiform gyrus (Supplementary Table 5). Evidence of MO-specific linear modulation in the OAS was limited to the superior parietal cortex, but also extended outside the OAS, in the lateral middle temporal gyrus (Supplementary Table 5). These two brain regions thus presented a concomitant subliminal (see main text) and linear activation profile as a function of Contrast, indicating significant activation both below and above PT, with relatively higher response amplitudes as the Contrast increased.

### fMRI activation as a function of PAS ratings (n = 12), conforming to the step BOLD amplitude model

In the inferior and superior parietal cortices, and in the lateral middle temporal gyrus, concomitantly to a subliminal BOLD modulation (see main text), we also found evidence of a step BOLD modulation (Supplementary Table 7), indicating significant activation both with and without SA, compared to the Empty baseline, but relatively stronger activation with than without SA. NO-specific step BOLD modulation was found in the bilateral fusiform gyri (Supplementary Table 7).

### fMRI activation as a function of Contrast (n = 24): Main effect of Contrast (increases)

We assessed whether the increase of image Contrast as a main effect, that is, independently of Manipulability level (MO, NO), induced an increase of brain activation. We found significant BOLD response increases along the visual pathways, including the posterior thalami, the striate and extra-striate occipital cortices, and the ventro-temporal cortex, extending medially to the amygdala. This largely symmetrical activation pattern, was complemented by the involvement, in the left hemisphere, of the anterior, inferior parietal cortex, the ventral premotor cortex, and the orbital inferior frontal cortex. A right-hemispheric activation in the pre- and post-central gyri was also found (Supplementary Fig. 4, Supplementary Table 9).

This BOLD response pattern is highly consistent with the well-characterized neural signature of conscious perception, with brain activity reverberating from the thalamus to sensory cortices, and from there to fronto-parietal cortices, giving rise to backward and forward circuit-level signal amplification^35,36^. This is in agreement with the idea that as image Contrast increased, the level of conscious perception of the stimulus also increased. Nested in this neural correlate of conscious perception, there may be a constellation of background executive processes, such as selective attention and self-monitoring^37^. This is however irrelevant in the context of the present study’s main focus, since in the content-specific comparisons between MO and NO, these additional confounding processes are mutually subtracted away.

---

**Supplementary Table 1.** Center coordinates for small volume Family Wise Error (smFWE) correction for multiple comparisons in the set of regions for tool-related cognition, as revealed by the meta-analysis of Ishibashi et al.^2^. Our neuroanatomical predictions were targeted to the premotor and parietal regions in this set, constituting the object-directed action representation system (OAS). Occipito-temporal regions, which in the same meta-analysis were identified as contributing to object identification alone, were also tested. ^§^Areas and cytoarchitectonic probabilities according to www.fz-juelich.de/ime/spm_anatomy_toolbox. L = left; R = right.

| Brain region (Brodmann area) [area, cytoarchitectonic probability]<sup>§</sup> | OAS | MNI coordinates (mm) |  |  |
| --- | --- | --- | --- | --- |
|  |  | x | y | z |
| L ventral premotor cortex (BA 6) | yes | -50.3 | 6.3 | 30.7 |
| L superior frontal gyrus / dorsal premotor cortex (BA 6) | yes | -20.6 | -4.4 | 62.2 |
| L inferior parietal cortex (BA 40) [area PFt, 64%] | yes | -45.7 | -31.7 | 44.8 |
| L superior parietal cortex (BA 7) [area 7A, 67%] | yes | -23.4 | -61.1 | 60.9 |
| L lateral middle temporal gyrus (BA 19/37) | no | -47.8 | -65.1 | -1.8 |
| L fusiform gyrus (BA 37) [area FG3, 83%] | no | -28.0 | -46.0 | -15.9 |
| R fusiform gyrus (BA 37) [area FG3, 85%] | no | 33.6 | -50.5 | -12.1 |

**Supplementary Table 2.** Statistical contrast weights for the analysis of BOLD activation as a function of Contrast level.

| Contrast name | MOThr<br>m2 -<br>Empty | MOThr<br>m1 -<br>Empty | MOThr0<br>- Empty | MOThrp<br>1 -<br>Empty | MOFull<br>- Empty | NOThr<br>m2 -<br>Empty | NOThr<br>m1 -<br>Empty | NOThr0<br>- Empty | NOThrp<br>1 -<br>Empty | NOFull -<br>Empty |
| --- | --- | --- | --- | --- | --- | --- | --- | --- | --- | --- |
| Simple effect of Manipulability for Contrast levels above PT (MO > NO) | 0 | 0 | 0 | +1 | +1 | 0 | 0 | 0 | -1 | -1 |
| Simple effect of Manipulability for Contrast levels above PT (NO > MO) | 0 | 0 | 0 | -1 | -1 | 0 | 0 | 0 | +1 | +1 |
| Main effect of Manipulability (MO > NO) | +1 | +1 | +1 | +1 | +1 | -1 | -1 | -1 | -1 | -1 |
| Main effect of Manipulability (NO > MO) | -1 | -1 | -1 | -1 | -1 | +1 | +1 | +1 | +1 | +1 |
| Main effect of Contrast (increases) | -2 | -1 | 0 | +1 | +2 | -2 | -1 | 0 | +1 | +2 |

b) First-level contrast weights, modeling either a subliminal, a step, or a linear effect.
| Contrast name | Empty | MOThr<br>m2 | MOThr<br>m1 | MOThr<br>0 | MOThr<br>p1 | MOFull | NOThr<br>m2 | NOThr<br>m1 | NOThr<br>0 | NOThr<br>p1 | NOFull |
| --- | --- | --- | --- | --- | --- | --- | --- | --- | --- | --- | --- |
| MO subliminal model | -5 | +1 | +1 | +1 | +1 | +1 | 0 | 0 | 0 | 0 | 0 |
| MO step model | -1 | -1 | -1 | -1 | +2 | +2 | 0 | 0 | 0 | 0 | 0 |
| MO linear model | -2.5 | -1.5 | -0.5 | +0.5 | +1.5 | +2.5 | 0 | 0 | 0 | 0 | 0 |
| NO subliminal model | -5 | 0 | 0 | 0 | 0 | 0 | +1 | +1 | +1 | +1 | +1 |
| NO step model | -1 | 0 | 0 | 0 | 0 | 0 | -1 | -1 | -1 | +2 | +2 |
| NO linear model | -2.5 | 0 | 0 | 0 | 0 | 0 | -1.5 | -0.5 | +0.5 | +1.5 | +2.5 |

**Supplementary Table 3.** Simple effect of Manipulability for image Contrast levels above PT (Thrp1, Full). MO-specific (MO > NO) and NO-specific (NO > MO) brain activation (n = 24). All reported effects passed a peak-level smFWE P < 0.05, except for the statistical trend (smFWE P < 0.1) in the ventral premotor cortex (marked by an asterisk). L = left; R = right. When, in addition to Contrast levels above PT (Thrp1, Full), we also considered those below (Thrm2, Thrm1) and at PT (Thr0), we found comparable effects (Main effect of Manipulability: Online Methods, Supplementary Table 4).

| Brain region | Voxels | P-value | Z(1,230) | MNI coordinates (mm) |  |  |
| --- | --- | --- | --- | --- | --- | --- |
|  |  |  |  | x | y | z |
| MO > NO |  |  |  |  |  |  |
| L ventral premotor cortex | 36 | 0.074* | 2.39 | -50 | 8 | 28 |
| L inferior parietal cortex | 11 | 0.021 | 2.93 | -50 | -30 | 40 |
| L superior parietal cortex | 4 | 0.022 | 2.90 | -28 | -60 | 60 |
| L lateral middle temporal gyrus | 23 | 0.002 | 3.72 | -50 | -70 | 4 |
| NO > MO |  |  |  |  |  |  |
| L fusiform gyrus | 22 | 0.001 | 3.94 | -24 | -44 | -12 |
| R fusiform gyrus | 22 | 4.5 x 10<sup>-8</sup> | 5.99 | 32 | -46 | -8 |

**Supplementary Table 4.** Main effect of Manipulability. MO-specific (MO > NO) and NO-specific (NO > MO) brain activation (n = 24). All reported effects passed a peak-level smFWE P < 0.05 correction. L = left; R = right.

| Brain region | Voxels | P-value | Z(1,230) | MNI coordinates (mm) |  |  |
| --- | --- | --- | --- | --- | --- | --- |
|  |  |  |  | x | y | z |
| MO > NO |  |  |  |  |  |  |
| L ventral premotor cortex | 17 | 0.016 | 3.03 | -50 | 8 | 28 |
| L inferior parietal cortex | 22 | 0.007 | 3.33 | -48 | -28 | 44 |
| L superior parietal cortex | 4 | 0.044 | 2.62 | -28 | -60 | 60 |
| L lateral middle temporal gyrus | 39 | 0.004 | 3.48 | -50 | -70 | -4 |
| NO > MO |  |  |  |  |  |  |
| L fusiform gyrus | 18 | 4.1 x 10<sup>-4</sup> | 4.15 | -26 | -42 | -12 |
| R fusiform gyrus | 21 | 8.5 x 10<sup>-9</sup> | 6.26 | 32 | -46 | -8 |

**Supplementary Table 5.** MO-specific (MO > NO) and NO-specific (NO > MO) brain activation as a function of Contrast (n = 24). All reported effects passed a peak-level smFWE P < 0.05 correction. L = left; R = right.

a) Step BOLD amplitude model
| Brain region | Voxels | P-value | Z(1,23) | MNI coordinates (mm) |  |  |
| --- | --- | --- | --- | --- | --- | --- |
|  |  |  |  | x | y | z |
| NO > MO |  |  |  |  |  |  |
| L fusiform gyrus | 27 | 0.047 | 2.67 | -26 | -46 | -12 |
| R fusiform gyrus | 22 | 0.004 | 3.62 | 30 | -48 | -8 |

b) Linear BOLD amplitude model
| Brain region | Voxels | P-value | Z(1,23) | MNI coordinates (mm) |  |  |
| --- | --- | --- | --- | --- | --- | --- |
|  |  |  |  | x | y | z |
| MO > NO |  |  |  |  |  |  |
| L superior parietal cortex | 2 | 0.026 | 2.92 | -28 | -62 | 60 |
| L lateral middle temporal gyrus | 16 | 0.016 | 3.10 | -50 | -70 | -4 |
| NO > MO |  |  |  |  |  |  |
| R fusiform gyrus | 12 | 0.001 | 3.89 | 30 | -48 | -8 |

**Supplementary Table 6.** Statistical contrast weights for the analysis of BOLD activation as a function of PAS ratings. First-level contrast weights, modeling either a subliminal effect (i.e. mean activation significant both with and without SA) or a step effect (i.e. mean activation significant only with SA).

| **Contrast name** | **Empty** | **MOPAS{1,2}** | **MOPAS{3,4}** | **NOPAS{1,2}** | **NOPAS{3,4}** |
| --- | --- | --- | --- | --- | --- |
| MO subliminal model | -2 | +1 | +1 | 0 | 0 |
| MO step model | -1 | -1 | +2 | 0 | 0 |
| NO subliminal model | -2 | 0 | 0 | +1 | +1 |
| NO step model | -1 | 0 | 0 | -1 | +2 |

**Supplementary Table 7.**
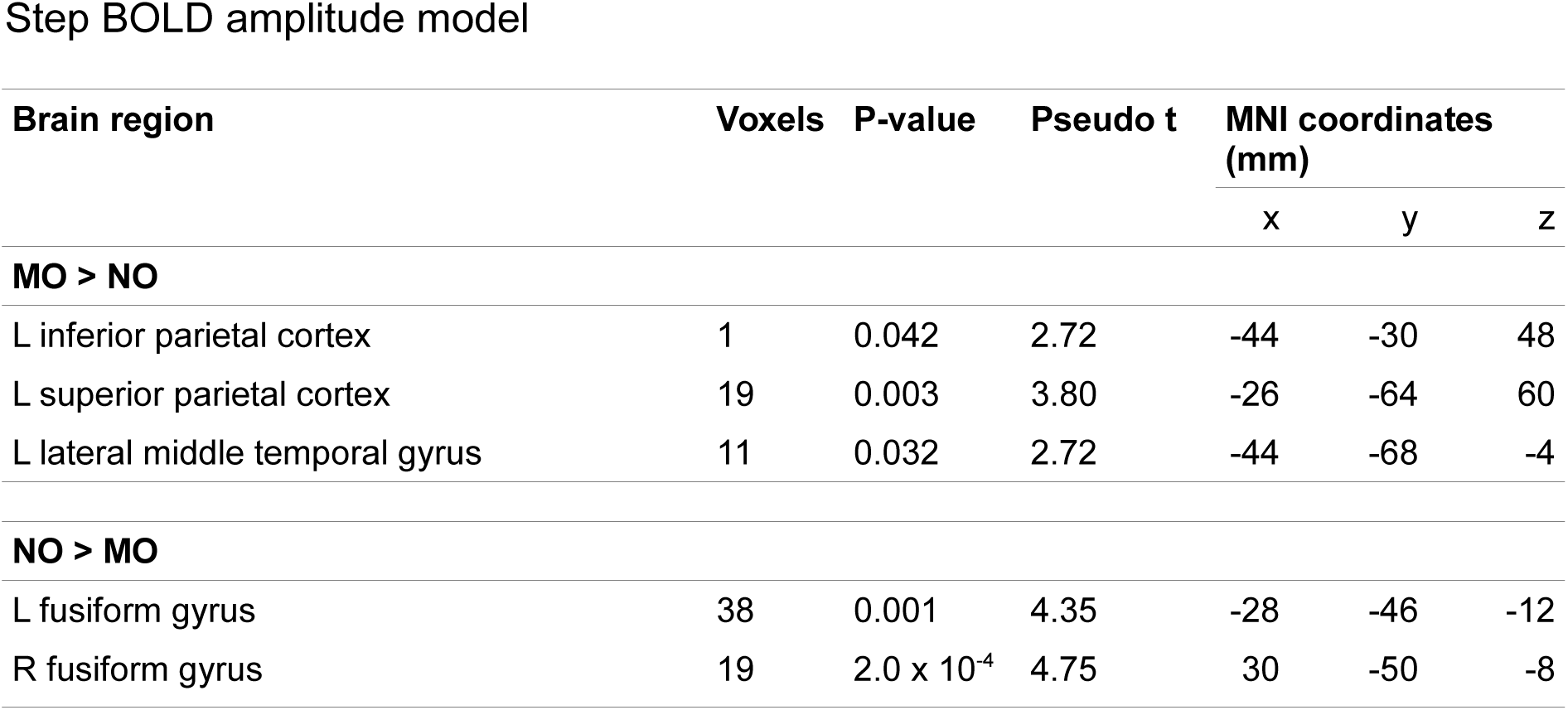
MO-specific (MO > NO) and NO-specific (NO > MO) brain activation as a function of PAS (n = 12). All reported effects passed a peak-level smFWE P < 0.05 correction. L = left; R = right.

**Supplementary Table 8.** MO-specific (MO > NO) and NO-specific (NO > MO) brain activation as a function of Contrast, in the sub-sample of 12 subjects included in the BOLD analysis as a function of PAS. All reported effects passed a peak-level smFWE P < 0.05 correction. L = left; R = right.

**a) Subliminal BOLD amplitude model**
| Brain region | Voxels | P-value | Pseudo t | MNI coordinates (mm) |  |  |
| --- | --- | --- | --- | --- | --- | --- |
|  |  |  |  | x | y | z |
| MO > NO |  |  |  |  |  |  |
| L ventral premotor cortex | 22 | 0.009 | 3.58 | -52 | 10 | 32 |
| L inferior parietal cortex | 27 | 0.009 | 3.68 | -48 | -28 | 44 |
| L superior parietal cortex | 17 | 0.007 | 3.67 | -22 | -66 | 60 |
| L lateral middle temporal gyrus | 29 | 0.009 | 3.45 | -46 | -64 | -4 |
| NO > MO |  |  |  |  |  |  |
| R fusiform gyrus | 14 | 0.002 | 4.37 | 30 | -50 | -8 |

**b) Step BOLD amplitude model**
| Brain region | Voxels | P-value | Pseudo t | MNI coordinates (mm) |  |  |
| --- | --- | --- | --- | --- | --- | --- |
|  |  |  |  | x | y | z |
| NO > MO |  |  |  |  |  |  |
| L fusiform gyrus | 23 | 0.011 | 3.32 | -28 | -46 | -12 |
| R fusiform gyrus | 12 | 0.006 | 3.92 | 30 | -50 | -8 |

**c) Linear BOLD amplitude model**
| Brain region | Voxels | P-value | Pseudo t | MNI coordinates (mm) |  |  |
| --- | --- | --- | --- | --- | --- | --- |
|  |  |  |  | x | y | z |
| MO > NO |  |  |  |  |  |  |
| L superior parietal cortex | 23 | 0.006 | 3.86 | -26 | -64 | 60 |
| L lateral middle temporal gyrus | 2 | 0.044 | 2.58 | -44 | -68 | -4 |
| NO > MO |  |  |  |  |  |  |
| L fusiform gyrus | 11 | 0.019 | 3.05 | -28 | -46 | -12 |
| R fusiform gyrus | 13 | 0.002 | 4.33 | 30 | -50 | -8 |

**Supplementary Table 9.** Main effect of Contrast (n = 24). All reported effects passed a peak-level P < 0.05, whole brain FWE correction for multiple comparisons. Indented brain regions belong to the same activation cluster as the non-indented brain region immediately above. ^§^Areas and cytoarchitectonic probabilities according to www.fz-juelich.de/ime/spm_anatomy_toolbox. L = left; R = right.

| Brain region [area, cytoarchitectonic probability]§ | Voxels | P-value | Z(1,230) | MNI coordinates (mm) |  |  |
| --- | --- | --- | --- | --- | --- | --- |
|  |  |  |  | x | y | z |
| BOLD increases with Contrast increase |  |  |  |  |  |  |
| L inferior frontal gyrus (pars orbitalis) | 48 | 2.0 x 10<sup>-4</sup> | 5.65 | -34 | 34 | 16 |
| L ventral premotor cortex | 9 | 0.010 | 4.86 | -44 | 6 | 32 |
| L inferior parietal cortex [area 2, 54%] | 4 | 0.041 | 4.52 | -44 | -38 | 48 |
| L inferior temporal gyrus | 1952 | 9.7 x 10<sup>-7</sup> | 6.56 | -40 | -16 | -32 |
| L fusiform gyrus [area FG3, 79%] | “ | < 1.0 x 10<sup>-9</sup> | 8.00 | -32 | -48 | -16 |
| L inferior occipital gyrus [area hOc4la, 63%] | “ | < 1.0 x 10<sup>-9</sup> | 8.00 | -42 | -76 | -8 |
| L middle occipital gyrus [area hOc4lp, 70%] | “ | < 1.0 x 10<sup>-9</sup> | 8.00 | -36 | -90 | 8 |
| L superior occipital gyrus | 18 | 0.002 | 5.16 | -26 | -70 | 36 |
| L thalamus [temporal, 58%] | 6 | 0.017 | 4.72 | -18 | -32 | 0 |
| L amygdala | 35 | 0.006 | 4.97 | -26 | -6 | -16 |
| R precentral gyrus | 211 | 2.7 x 10<sup>-7</sup> | 6.76 | 44 | -14 | 52 |
| R postcentral gyrus | “ | 2.8 x 10<sup>-7</sup> | 6.75 | 46 | -20 | 48 |
| R inferior temporal gyrus | 2378 | 1.0 x 10<sup>-5</sup> | 6.13 | 38 | -12 | -36 |
| R fusiform gyrus [area FG3, 91%] | “ | < 1.0 x 10<sup>-9</sup> | 8.00 | 30 | -44 | -16 |
| R lingual gyrus [area hOc3v, 56%] | “ | 0.001 | 5.43 | 24 | -92 | -8 |
| R calcarine gyrus [area hOc1, 76%] | “ | 0.010 | 4.84 | 18 | -98 | -4 |
| R inferior occipital gyrus [area hOc4lp, 68%] | “ | < 1.0 x 10<sup>-9</sup> | 8.00 | 38 | -88 | -4 |
| R inferior occipital gyrus [area hOc1, 51%] | “ | 2.0 x 10<sup>-4</sup> | 5.64 | 24 | -98 | -4 |
| R middle occipital gyrus | “ | 2.1 x 10<sup>-8</sup> | 7.14 | 34 | -82 | 20 |
| R superior occipital gyrus | “ | 7.0 x 10<sup>-4</sup> | 5.54 | 28 | -74 | 36 |
| R thalamus [temporal, 54%] | “ | 2.0 x 10<sup>-5</sup> | 6.07 | 20 | -32 | 0 |
| R amygdala | 8 | 0.009 | 4.87 | 26 | -4 | -16 |

